# Regulating the proinflammatory response to implanted composite biomaterials comprising polylactide and hydroxyapatite by targeting immunometabolism

**DOI:** 10.1101/2023.10.21.563447

**Authors:** Chima V. Maduka, Ashley V. Makela, Evran Ural, Katlin B. Stivers, Maxwell M. Kuhnert, Anthony Tundo, Mohammed Alhaj, Ehsanul Hoque Apu, Nureddin Ashammakhi, Kurt D. Hankenson, Ramani Narayan, Jennifer H. Elisseeff, Christopher H. Contag

## Abstract

Composite biomaterials comprising polylactide (PLA) and hydroxyapatite (HA) are applied in bone, cartilage and dental regenerative medicine, where HA confers osteoconductive properties. However, after surgical implantation, adverse immune responses to these composites can occur, which have been attributed to size and morphology of HA particles. Approaches to effectively modulate these adverse immune responses have not been described. PLA degradation products have been shown to alter immune cell metabolism, which drives the inflammatory response. Therefore, we aimed to modulate the inflammatory response to composite biomaterials by regulating glycolytic flux with small molecule inhibitors incorporated into composites comprised of amorphous PLA (aPLA) and HA (aPLA+HA). Inhibition at specific steps in glycolysis reduced proinflammatory (CD86^+^CD206^-^) and increased pro-regenerative (CD206^+^) immune cell populations around implanted aPLA+HA resulting in a pro-regenerative microenvironment. Notably, neutrophil and dendritic cell (DC) numbers along with proinflammatory monocyte and macrophage populations were decreased, and Arginase 1 expression among DCs was increased. Targeting immunometabolism to control the inflammatory response to biomaterial composites, and creating a pro-regenerative microenvironment, is a significant advance in tissue engineering where immunomodulation enhances osseointegration, and angiogenesis, which will lead to improved bone regeneration.

## Introduction

Composite biomaterials comprising polylactide (PLA) and hydroxyapatite (HA) are often fabricated for clinical applications involving bone, cartilage and dental regenerative engineering. The combination of PLA with HA overcomes the brittleness of HA, while increasing the tensile modulus and hardness of the polymer to approximate that of trabecular bone^1-5^. Importantly, because 70% (by weight)^6,7^ of bone tissue is HA, HA is bioactive and can facilitate new bone^8-10^, cartilage^11^ and dental^12^ tissue formation. Thus, HA confers osteoconductive characteristics to composite biomaterial formulations. Accordingly, it has been shown that inclusion of, or coating with, HA enhances bone-implant integration (osseointegration)^13-15^. In turn, enhanced osseointegration has been shown to reduce the risk of implant failure, and thus increase the longevity of total knee and hip replacements^16-19^.

Hydrolytic byproducts of PLA degradation were previously thought to elicit the foreign body response by reduced pH in the biomaterial microenvironment^20^. This notion originated from the correlation of decreased bioluminescence of the bacterium *Photobacterium phosphoreum* with reduced pH in unbuffered solutions containing PLA breakdown products^21^. However, recent advances now demonstrate that hydrolytic byproducts of PLA degradation activate surrounding immune cells by altering cellular bioenergetics and significantly increasing glycolytic flux^22,23^. In a manner dependent on CCR2 and CX3CR1 signaling, immunometabolic cues regulate the trafficking of circulating monocytes to the PLA biomaterial microenvironment^24^. Consequently, targeting metabolic reprogramming effectively modulated the foreign body response to implanted PLA as demonstrated by reduced neutrophil recruitment, increased IL-4 production from T helper 2 cells and γδ+ T-cells, and skewing of monocyte, macrophage and dendritic cell populations toward pro-regenerative phenotypes^22,24^.

Compared to PLA alone, incorporation of HA exerts immunomodulatory effects by: a) decreasing the relative levels of proinflammatory (CD86^+^CD206^-^) dendritic cells (DCs), including proinflammatory DCs expressing class II major histocompatibility complex (MHC II); b) increasing the relative levels of transition (CD86^+^CD206^+^) and anti-inflammatory or pro-regenerative (CD206^+^) DCs, including those expressing MHC II; c) reducing the relative levels of proinflammatory monocytes and macrophages relative to transition cell populations in the biomaterial microenvironment^24^. Collectively, these immunomodulatory effects are able to enhance osseointegration and angiogenesis during skeletal tissue regeneration^9,14^, with transition immune cells playing a crucial role^25,26^. However, HA alone^13,27-29^ or as a composite with PLA^24,30^ chronically activates neutrophils counteracting its beneficial immunological effects. Accordingly, short-term studies^31,32^ are more likely to report beneficial immunomodulatory effects than long-term (> 2 years) studies^33^ when applying bulk composite implants comprising HA and PLA. Moreover, wear particles of HA (from coated metal implants applied in total knee and hip replacements) are known to drive chronic inflammation leading to implant failure^13,15^, a process that is affected by particle size^29,34,35^ and morphology^36^. In fact, HA wear particles have been shown to be recognized by the opsonin receptor^37^ as well as by Toll-like receptor 4^15,38^, upregulating proinflammatory genes^39^, and activating both nuclear factor-kappa B and interferon regulatory factor 3^38^ to trigger the production of proinflammatory cytokines in a manner dependent on the membrane proximal kinase, *Syk*, as well as members of the mitogen-activated protein kinase family of signaling molecules^40^. In the bone microenvironment, this could drive osteolysis^33^ by increasing osteoclastogenesis through upregulating M-CSF and RANKL^41^.

Despite aforementioned in-vitro and in-vivo evidence that adverse immune responses could occur with HA and composites of HA with PLA, corresponding immunomodulatory strategies are yet to be explored. Furthermore, while the majority of prior studies have focused on the inflammatory underpinnings of HA particles^15,29,34,35,38,40,42^, developing immunomodulatory strategies for bulk HA composites have not been addressed. Here, we demonstrate that modulating glycolytic flux by incorporating small molecule inhibitors of glycolysis into a composite material comprised of amorphous polylactide (aPLA) and HA (aPLA+HA) leads to a pro-regenerative implant microenvironment. We observed that inhibiting different glycolytic steps reduces the proportion of proinflammatory signals and increases the relative levels of pro-regenerative CD45^+^ immune cell population in the microenvironment surrounding implanted PLA-HA composite biomaterials. Notably, Ly6G^+^ neutrophil populations are decreased following incorporation of glycolytic inhibitors in aPLA+HA composites. While overall levels of recruited CD11b^+^ monocytes and F4/80^+^ tissue macrophages do not decrease with the incorporation of glycolytic inhibitors in aPLA+HA composites, the respective proinflammatory proportions of these populations are reduced. In addition, dendritic cell populations are reduced by the incorporation of glycolytic inhibitors in implanted PLA-HA composites. Notably, Arginase 1 levels were increased in dendritic cells and dendritic cells expressing MHC II by incorporation of a glycolytic inhibitor. Control of inflammatory responses to biomaterial composites using glycolytic inhibitors is a significant advancement that could lead to enhanced osseointegration and angiogenesis by generating a pro-regenerative microenvironment resulting in improved tissue regeneration^9,14,43^.

## Result and Discussion

Having observed the pivotal role that altered metabolism plays in the foreign body response to polylactide^22-24^, we hypothesized that locally modifying immunometabolic cues in the composite biomaterial microenvironment will modulate adverse inflammatory responses. To test this hypothesis, we subcutaneously implanted aPLA+HA biomaterials, with or without incorporated aminooxyacetic acid (a.a.) or 2-deoxyglucose (2DG) at previously optimized doses,^22-24^ into C57BL/6J mice. These two small molecule inhibitors act at different steps in glycolysis; a.a. inhibits uptake of glycolytic substrates and glutamine metabolism, 2DG inhibits hexokinase in the glycolytic pathway^44,45^. For reference, sham controls where incisions were made without biomaterial implantation were included.

Although implantation of aPLA+HA increased overall nucleated hematopoietic (CD45^+^) cell populations, incorporation of a.a. but not 2DG reduced CD45^+^ levels (Fig. 1a-e). To understand the relative levels of polarized CD45^+^ populations, we designated proinflammatory subsets^46^ as CD86^+^CD206^-^ and anti-inflammatory subsets^43^ as CD206^+^. Relative to sham controls, the fold change of proinflammatory CD45^+^ cells with respect to anti-inflammatory CD45^+^ cells was elevated (Fig. 1f). However, incorporation of either a.a. or 2DG reduced proinflammatory CD45^+^ proportions in comparison to aPLA+HA alone (Fig. 1f). Interestingly, implantation of aPLA+HA decreased the fold change of anti-inflammatory CD45^+^ cells with respect to proinflammatory CD45^+^ cells, likely due to the polylactide content^24^ of the composite biomaterial. In contrast, incorporation of either a.a. or 2DG increased anti-inflammatory CD45^+^ levels compared to aPLA+HA only (Fig. 1g-k). Moreover, incorporating a.a. tended to increase the frequency of CD45^+^ cells expressing Arginase 1 (Arg 1^+^) even though this trend was not statistically significant (Fig. 1l).

**Figure 1.**
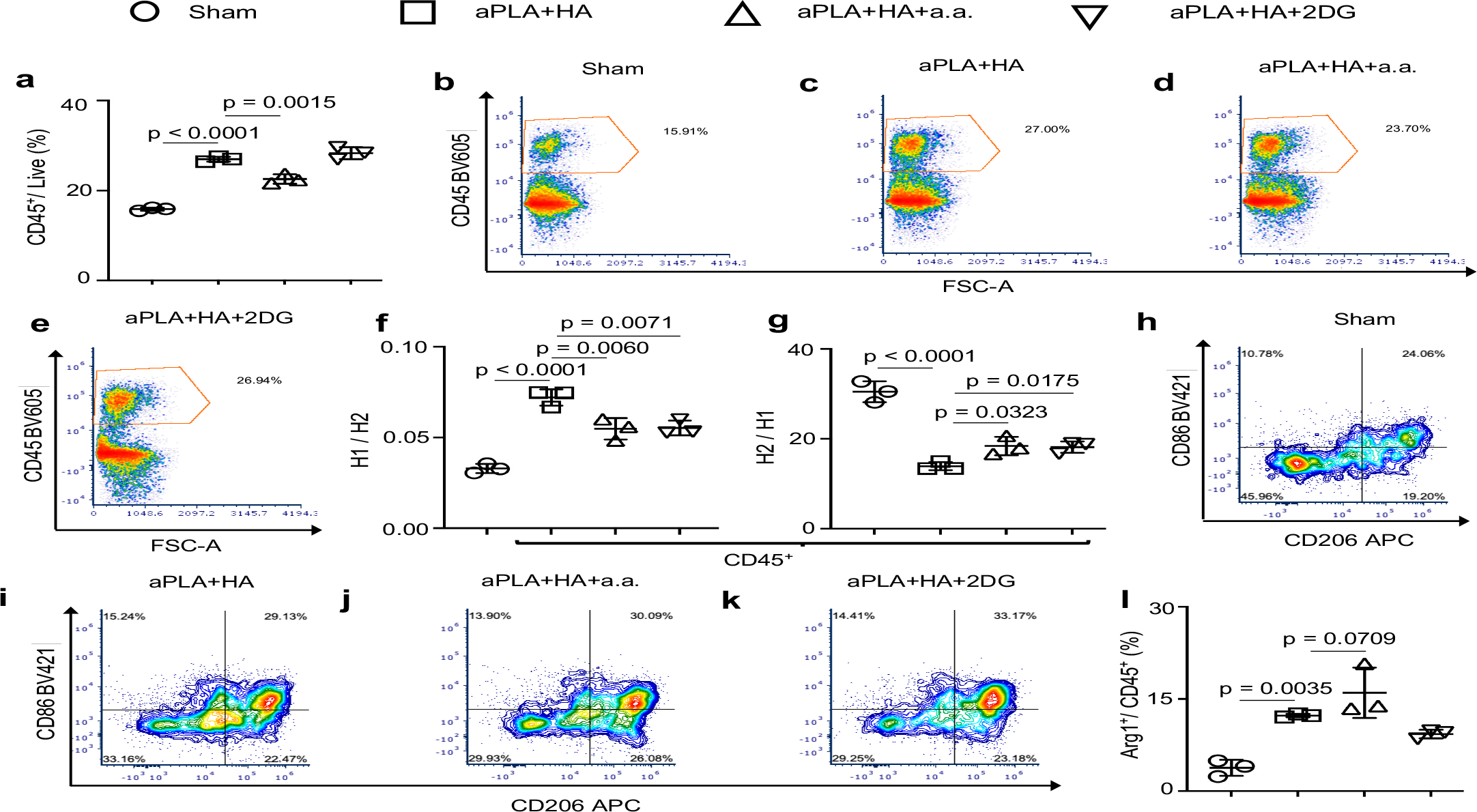
Glycolytic inhibition in the amorphous polylactide-hydroxyapatite composite biomaterial microenvironment modifies the numbers and inflammatory states of recruited nucleated hematopoietic cell populations. a-e, Flow cytometry quantification (a) and representative plots (b-e) of nucleated hematopoietic (CD45^+^) cells gated on live cells. f, Fold change of proinflammatory (H1; CD86^+^CD206^-^) cells with respect to anti-inflammatory (H2; CD206^+^) cells. g, Fold change of H2 with respect to H1 cells. h-k, Representative flow plots of CD86 and CD206 cells gated on CD45^+^ cells. l, Quantification of Arginase 1 (Arg1^+^) cells gated on CD45^+^ populations. One-way ANOVA followed by Tukey’s or Newman-Keul’s multiple comparison test, n = 3; amorphous polylactide (aPLA), hydroxyapatite (HA), aminooxyacetic acid (a.a.), 2-deoxyglucose (2DG).

Previously, we have observed that, relative to aPLA alone, aPLA+HA does not reduce Ly6G^+^ neutrophils recruited to the biomaterial microenvironment^24^. Here, we found that, compared to sham controls, aPLA+HA implantation elevated neutrophil levels (Fig. 2a-e). Remarkably, incorporating a.a. or 2DG in aPLA+HA modulated this proinflammatory tendency (Fig. 2a-e). Elevated neutrophil levels are prevalent in murine bone defects implanted with micron-sized HA particles, an effect that is reduced by using nano-sized HA particles^29^. Reduced neutrophil levels are correlated with the pro-regenerative macrophage phenotype that is necessary to drive bone regeneration^29^. This observation is translationally relevant as HA potently activates human neutrophils in-vitro^13,27,28^.

**Figure 2.**
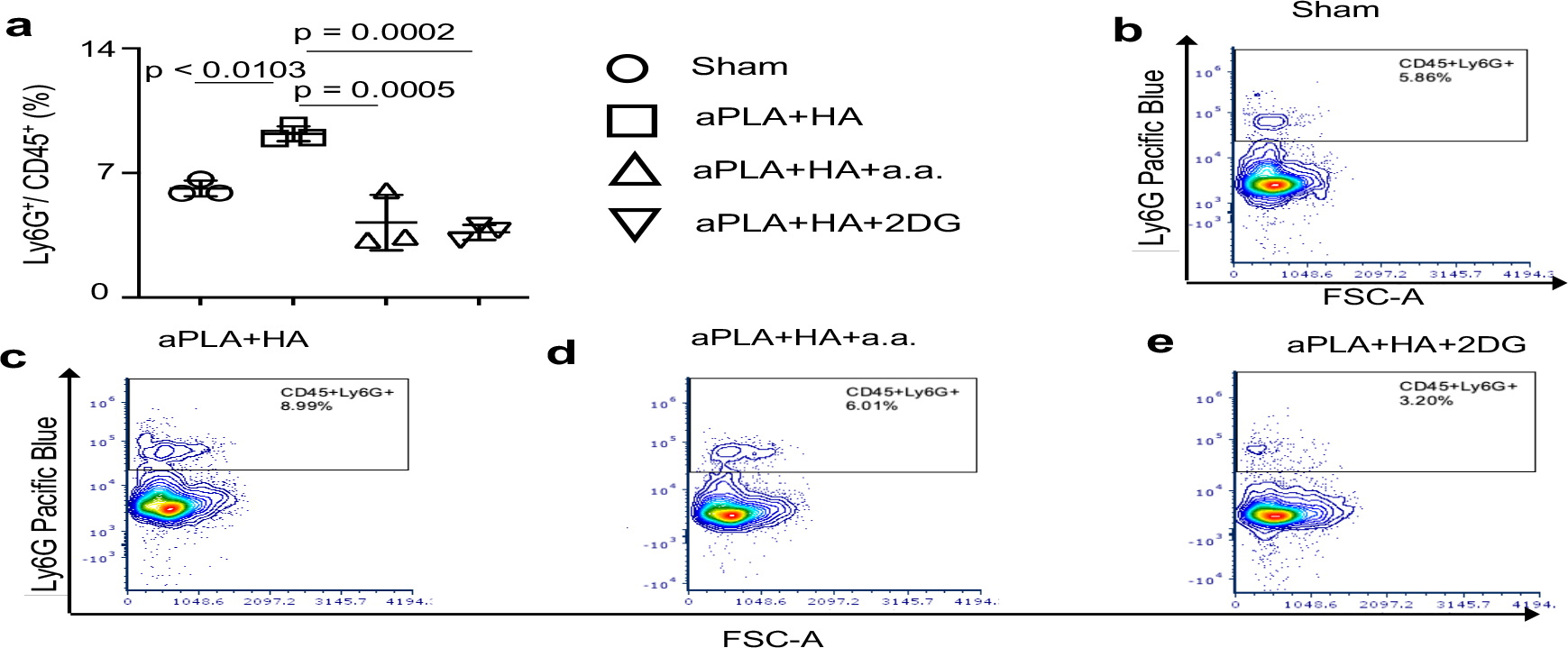
Incorporation of metabolic inhibitors modulate neutrophil recruitment in the amorphous polylactide-hydroxyapatite composite biomaterial microenvironment. a-e, Quantification (a) and representative flow cytometry plots (b-e) of neutrophils (Ly6G^+^) cells gated on CD45^+^ populations. One-way ANOVA followed by Tukey’s multiple comparison test, n = 3; amorphous polylactide (aPLA), hydroxyapatite (HA), aminooxyacetic acid (a.a.), 2-deoxyglucose (2DG).

Next, we examined the levels of CD11b^+^ monocytes^47-49^ and F4/80^+^ macrophages^50^ in the biomaterial microenvironment (CD11b is also expressed on some B-cells, neutrophils and macrophages^50^). Consistent with prior observations^30,51^, implantation of aPLA+HA increased monocyte and macrophage proportions relative to sham controls, but incorporation of a.a. or 2DG did not reduce cellular recruitment (Fig. 3a-f). We observed that aPLA+HA increased Arg1 levels among monocytes (Fig. 3g) and macrophages (Fig. 3h) relative to sham controls, likely from its immunomodulatory capability^9,29^. Additionally, incorporation of a.a. but not 2DG to aPLA+HA tended to further increase Arg1 levels among monocytes and macrophages, although this trend was not statistically significant (Fig. 3g-h). Relative to sham controls, aPLA+HA increased the fold change of proinflammatory monocytes with respect to anti-inflammatory monocytes; however, incorporation of either a.a. or 2DG reduced proinflammatory levels (Fig. 4a). Although implantation of aPLA+HA decreased the fold change of anti-inflammatory monocytes to proinflammatory monocytes, the tendency for a.a. to increase anti-inflammatory proportions was not statistically significant (Fig. 4b-f). Similar to observations made with monocytes, aPLA+HA elevated proinflammatory and reduced anti-inflammatory macrophage levels compared to sham controls (Fig. 4g-j). While incorporation of either a.a. or 2DG reduced proinflammatory macrophage levels, anti-inflammatory levels were not increased as observed in quantitated data and representative dot plots (Fig. 4g-l).

**Figure 3.**
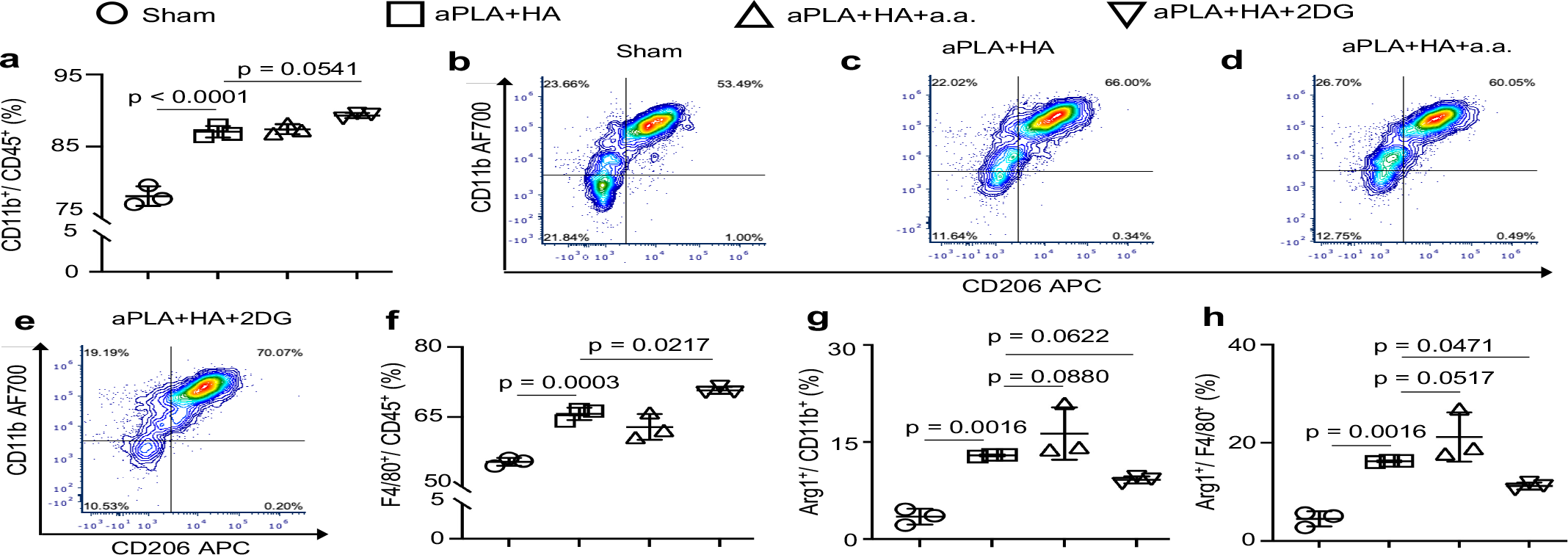
Monocyte and macrophage populations are differentially affected by targeting different glycolytic steps in the implanted amorphous polylactide-hydroxyapatite composite biomaterial microenvironment. a, Flow cytometry quantification of monocytes (CD11b^+^ cells) gated on CD45^+^ populations. b-e, Representative plots of monocytes (CD11b^+^) and macrophages (F4/80^+^) gated on CD45^+^ populations. f, Quantification of macrophages in the composite biomaterial microenvironment. g-h, Quantification of Arginase 1 (Arg1^+^) monocytes (g) and macrophages (h). One-way ANOVA followed by Tukey’s or Newman-Keul’s multiple comparison test, n = 3; amorphous polylactide (aPLA), hydroxyapatite (HA), aminooxyacetic acid (a.a.), 2-deoxyglucose (2DG).

**Figure 4.**
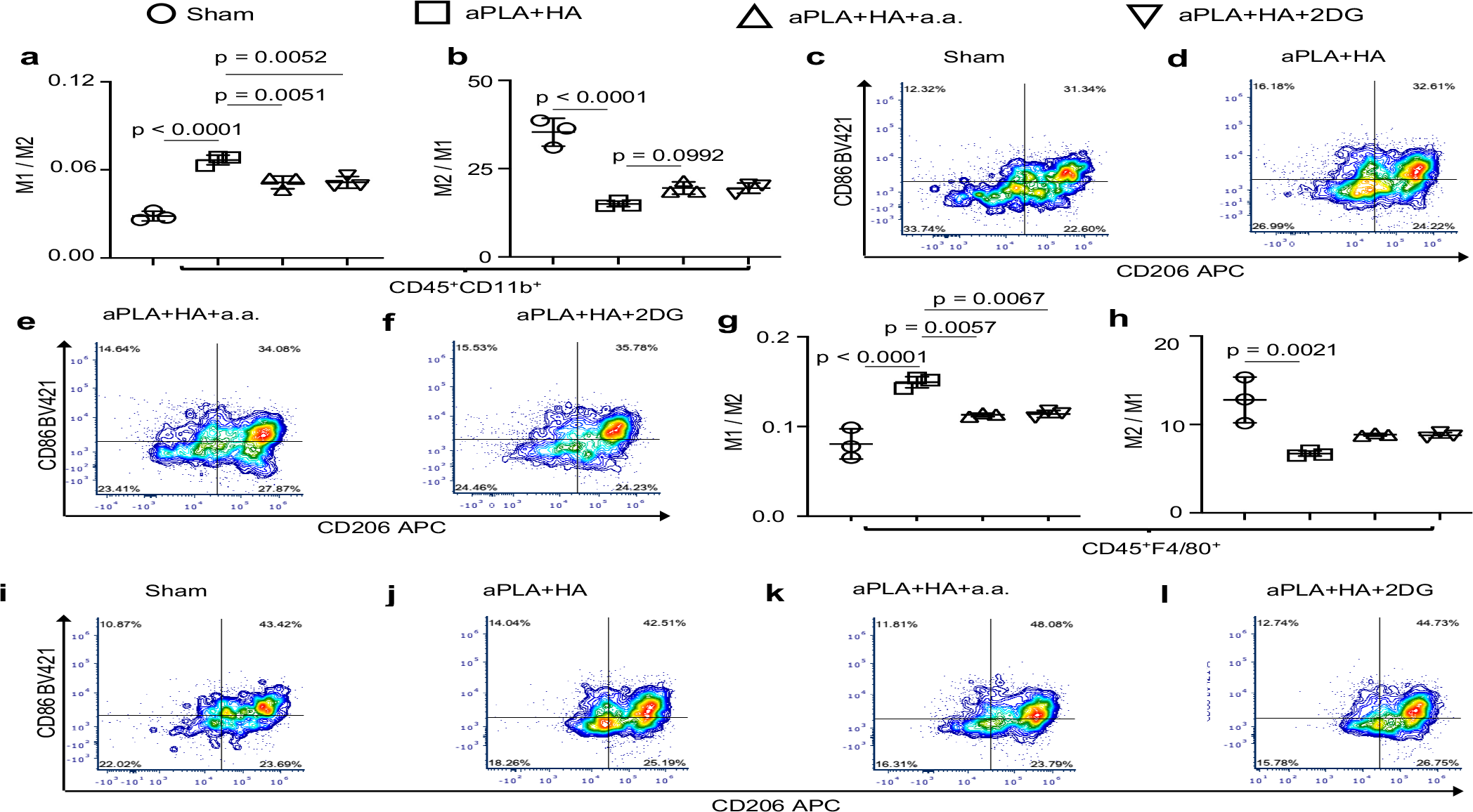
Activation states of monocytes and macrophages are modulated by glycolytic inhibition in the amorphous polylactide-hydroxyapatite composite biomaterial microenvironment. a, Fold change of proinflammatory (M1; CD86^+^CD206^-^) monocytes with respect to anti-inflammatory (M2; CD206^+^) monocytes. b, Fold change of M2 monocytes with respect to M1 monocytes. c-f, Representative plots of CD86 and CD206 cells gated on monocyte populations (CD45^+^CD11b^+^). g, Fold change of proinflammatory (M1; CD86^+^CD206^-^) macrophages with respect to anti-inflammatory (M2; CD206^+^) macrophages. h, Fold change of M2 macrophages with respect to M1 macrophages. i-l, Representative plots of CD86 and CD206 cells gated on macrophage populations (CD45^+^F4/80b^+^). One-way ANOVA followed by Tukey’s multiple comparison test, n = 3; amorphous polylactide (aPLA), hydroxyapatite (HA), aminooxyacetic acid (a.a.), 2-deoxyglucose (2DG).

We observed that CD11c^+^ dendritic cell populations were elevated in the aPLA+HA microenvironment compared to sham controls as previously reported^30,51^, and that incorporation of either a.a. or 2DG reduced these dendritic cell numbers (Fig. 5a-e). Interestingly, compared to sham controls, the fold change of proinflammatory dendritic cells relative to anti-inflammatory dendritic cells was increased in the microenvironment of aPLA+HA implants; yet, incorporation of a.a. or 2DG did not reduce proinflammatory dendritic cell levels (Fig. 5f). Furthermore, although the fold change of anti-inflammatory dendritic cells to proinflammatory dendritic cells was decreased in aPLA+HA compared to sham controls, incorporating a.a. or 2DG did not increase anti-inflammatory dendritic cells proportions (Fig. 5g).

**Figure 5.**
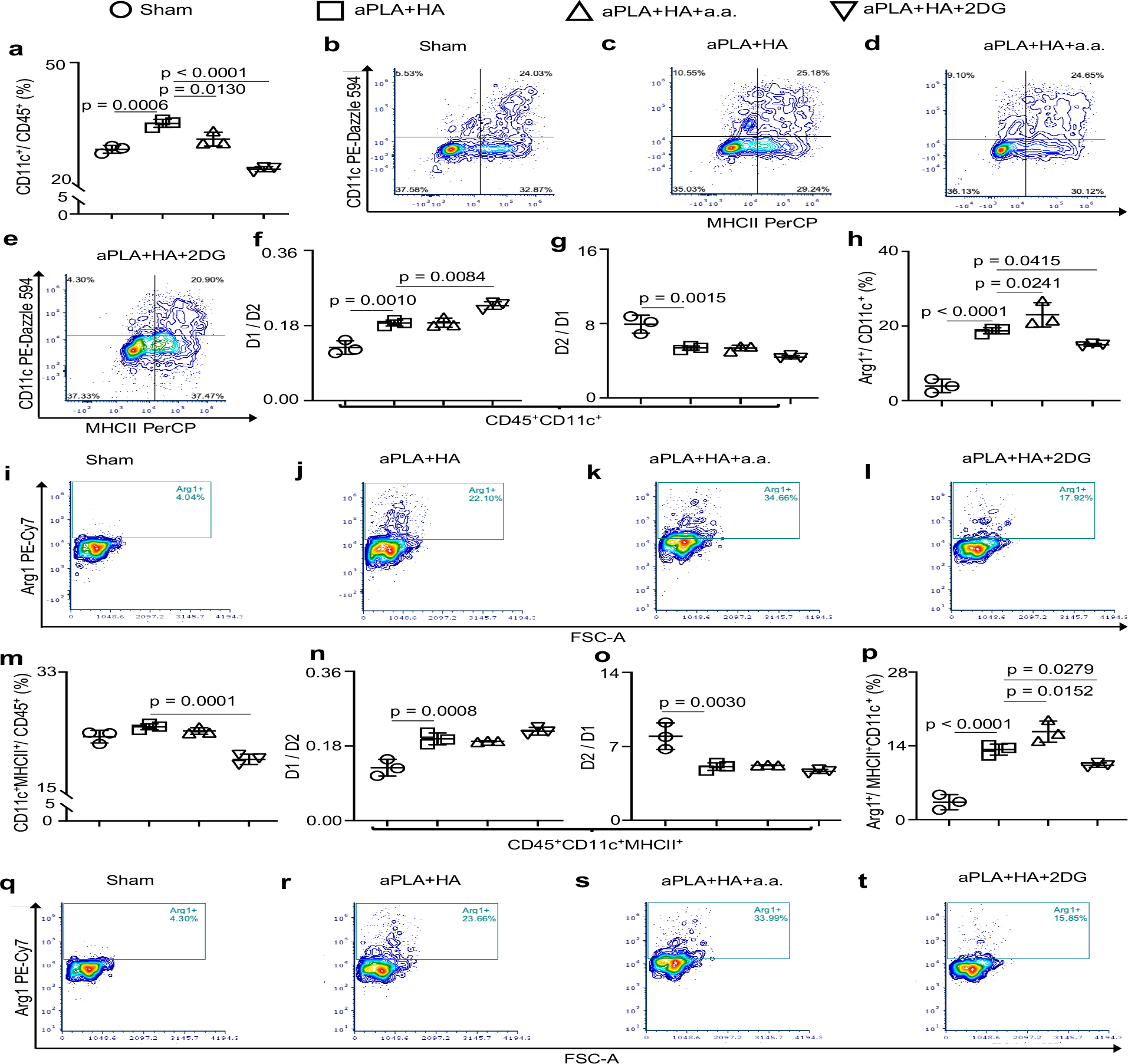
Proportions and inflammatory states of dendritic cells are affected in the amorphous polylactide-hydroxyapatite composite biomaterial microenvironment. a, Flow cytometry quantification of dendritic (CD11c^+^) cells gated on CD45^+^ cells. b-e, Representative plots of dendritic (CD11c^+^) cells with and without MHCII gated on CD45^+^ cells. f, Fold change of proinflammatory (D1; CD86^+^CD206^-^) dendritic cells with respect to anti-inflammatory (D2; CD206^+^) dendritic cells. g, Fold change of D2 with respect to D1 dendritic cells. h-l, Quantification (h) and representative plots (i-l) of Arginase 1 (Arg1^+^) dendritic cells. m, Dendritic cells expressing MHCII gated on CD45^+^ cells. n, Fold change of proinflammatory (D1; CD86^+^CD206^-^) dendritic cells expressing MHCII with respect to anti-inflammatory (D2; CD206^+^) dendritic cells expressing MHCII. o, Fold change of D2 with respect to D1 dendritic cells expressing MHCII. p-t, Quantification (p) and representative plots (q-t) of Arginase 1 (Arg1^+^) dendritic cells expressing MHCII. One-way ANOVA followed by Tukey’s or Newman-Keul’s multiple comparison test, n = 3; amorphous polylactide (aPLA), hydroxyapatite (HA), aminooxyacetic acid (a.a.), 2-deoxyglucose (2DG).

Expression of Arg1 among dendritic cells was increased following implantation of aPLA+HA relative to sham controls (Fig. 5h-j). Notably, compared to aPLA+HA, incorporating a.a. further elevated Arg1 expression among dendritic cell populations (Fig. 5h-l). Dendritic cells expressing MHC II were similar between sham and aPLA+HA groups; incorporating 2DG decreased the numbers of dendritic cells expressing MHC II when compared to aPLA+HA (Fig. 5m). In comparison to sham controls, aPLA+HA both increased proinflammatory and decreased anti-inflammatory proportions of dendritic cells expressing MHC II (Fig. 5n-o), revealing previously unappreciated mechanisms by which composite biomaterials drive proinflammatory states. Compared to aPLA+HA only, incorporation of a.a. elevated Arg1 expression among dendritic cells expressing MHC II (Fig. 5p-t). Increased Arg1 expression in the composite biomaterial microenvironment could arise from inhibition of aspartate-aminotransferase by a.a., which obviates metabolic and transcriptional activation of immune cells into proinflammatory states^52^. Elevated Arg1 is a crucial driver of osteoinduction, creating a pro-regenerative composite biomaterial microenvironment^53^.

In conclusion, we uncover new ways by which composite biomaterials affect the immune microenvironment, such as altering the ratio of proinflammatory to anti-inflammatory skewed CD45^+^ populations. We demonstrate that controlling metabolic states by modifying glycolytic flux around implanted composite biomaterials is capable of: a) decreasing neutrophil recruitment; b) decreasing proinflammatory monocyte and macrophage populations; c) decreasing dendritic cell numbers; d) and increasing Arg1expression among dendritic cells and dendritic cells expressing MHC II. Aminoxyacetic acid (a.a.), one of the metabolic inhibitors, has already been used safely in clinical trials for the treatment of other disease conditions^54^, making it a translatable small molecule for incorporation into composite biomaterials for future clinical use. Prior to translation, additional studies are needed to characterize the release profiles of metabolic inhibitors from composite biomaterials as well as the effects of implanting composite biomaterials containing embedded metabolic inhibitors in musculoskeletal tissues, such as bone defects, for regenerative medicine applications.

## Materials and Methods

### Metabolic inhibitors and their incorporation into biomaterials

The metabolic modulators 2-deoxyglucose (2DG; MilliporeSigma) and aminooxyacetic acid hemihydrochloride (a.a.; Sigma-Aldrich) were incorporated into composite biomaterials comprising amorphous polylactide (aPLA; PLA 4060D) and hydroxyapatite (HA; 2.5 μm particle sizes; Sigma-Aldrich) by melt-blending at 190°C for 3 min in a DSM 15 cc mini-extruder (DSM Xplore). Based on prior in-vitro studies^22^, 189 mg of 2DG, 90 mg of a.a. and 200 mg of HA were compounded in 10 g of aPLA to approximate effective in-vitro concentrations. Following extrusion from the DSM, a pelletizer (Leistritz Extrusion Technology) was used to create pellets. A second extrusion (Filabot EX2; 170°C with air set at 93) into 1.75 mm diameter filaments was undertaken. Filaments were cut into 1 mm long sizes for implantation into mice and sterilized for 30 min by ultraviolet radiation.

### Mouse model

Female C57BL/6J mice (9 weeks old) were obtained from Jackson Laboratory. Animal studies were approved by the Institutional Animal Care and Use Committee at Michigan State University (PROTO202100327). For the implantation of biomaterials, mice were anesthetized using 2-3% isofluorane mixed with oxygen. At the site of subcutaneous surgical implantation, the fur was shaved followed by disinfection of the skin using iodine and alcohol swabs. A pouch was made in the subcutis, followed by implantation of 1 mm long filaments of composite materials comprised of amorphous polylactide (aPLA) and HA (aPLA+HA), with and without incorporating a.a. and 2DG. Also included was a sham group with incision on the back of mice and pouch creation without biomaterial implantation. In all cases, surgical glue (3M Vetbond) was used to close the skin. Mice then received intraperitoneal or subcutaneous meloxicam (5 mg/kg) analgesia as well as postoperative saline.

### Tissue digestion protocol

Eleven weeks following implantation, mice were shaved around the implanted biomaterial site (or sham site), then euthanized for excision of tissue. Circular biopsies (8 mm diameter) were collected from each mouse and tissues were pooled from the same groups. Tissues were cut with surgical scissors for ∼1 min followed by digestion in an enzyme cocktail containing 0.5 mg/mL Liberase (Sigma-Aldrich), 0.5 mg/mL Collagenase Type IV (Stem Cell Technologies), 250 U/mL Deoxyribonuclease I (Worthington Biochemical Corporation) in 25 mM HEPES buffer (Sigma-Aldrich). The tissue/enzyme cocktail was incubated at 37°C with 5% CO2 on top of an orbital shaker, shaking at 70 rpm for 1 hour. Following incubation, 5 mL of the tissue/enzyme cocktail mixture was run through a 70 μm filter into a 50 mL conical tube and the remaining tissues which were not digested were mechanically dissociated against the serrated portion of a petri dish. The resultant mixture was filtered into the previous 50 mL conical tube. Remaining undigested tissue in the 70 μm filter was again mechanically dissociated with the thumb press of a syringe plunger for optimal extraction of cells. The petri dish was washed with cold Hanks’ Balanced Salt Solution without calcium, magnesium and phenol red (ThermoFisher Scientific), followed by filtration into the conical tube. Cells were centrifuged at 350G for 10 mins followed by automated counting (Countess Automated Cell Counter, Invitrogen) for flow cytometry.

### Flow cytometry

We used 1x10^6^ cells per sample for flow cytometry staining in a polypropylene 96-well round bottom plate (Sigma, cat#P6866). All staining steps were performed in 100 μL volume in the dark at 4°C. Samples were first incubated with LIVE/ DEAD Fixable Blue Dead Cell Stain kit (1:500, Thermofisher, cat#L23105) for 20 min. Thereafter, cells were washed once with flow buffer, followed by incubation with TruStain FcX (anti-mouse CD16/32) Antibody (BioLegend, Cat#101319; 1 μg/sample) in 50 μL volume for 10 min. The following antibodies were mixed together at 2x concentration in 50 μL and added directly to the cell suspension: BV605 CD45 (1:500, Biolegend, cat#103139), AF700 CD11b (1:300, Biolegend, cat#101222), BV785 F4/80 (1:300, Biolegend, cat#123141), BV421 CD86 (1:200, Biolegend, cat#105031), APC CD206 (1:200, Biolegend, cat#141707), PerCP MHCII (1:200, Biolegend, cat#107623), PacBlue Ly6G (1:250, BD Bioscience, cat#127611) and PE-Dazzle 594 CD11c (1:500, Biolegend, cat#117347). Cells and antibody mixture were incubated for 30 min. Cells were washed once prior to fixation and permeabilization (BD Cytofix/Cytoperm kit, cat#BDB554714) as per manufacturer’s instructions. Cells were then resuspended in BD Perm/wash buffer with PE-Cy7 Arg1 (1:100, ThermoFisher, cat#25-3697-80). Cells were incubated with antibody mixture for 30 min. Cells were washed twice with BD Perm/wash buffer followed by resuspension in a final volume of 100 μL for flow cytometry analysis.

The Cytek Aurora spectral flow cytometer (Cytek Biosciences, CA, USA) was used to analyze samples. Furthermore, fluorescence minus one (FMO) samples guided gating strategies, and the software FCSExpress (DeNovo Software, CA, USA) was used to analyze flow cytometry data.

### Statistics

We used GraphPad Prism as our software (GraphPad Prism Version 9.5.1) for statistical data analysis. Results are presented as mean ± standard deviation (SD), with figure legends showing exact statistical test, p-values and sample sizes.

## Statistics and reproducibility

We used GraphPad Prism Version 9.5.1 (528)) to analyse data presented as mean with standard deviation (SD), and provided information on statistical test, p-values and sample sizes in respective figure legends.

## Data availability

The data supporting the findings of this study are available within the paper and its Supporting Information.

## Acknowledgements

Funding for this work was provided in part by the James and Kathleen Cornelius Endowment at MSU.

## Author contributions

Conceptualization, C.V.M. and C.H.C.; Methodology, C.V.M., A.V.M., E.U., K.B.S., M.M.K, A.T., M.A., E.H.A, N.A., K.D.H., R.N., J.H.E., and C.H.C.; Investigation, C.V.M., A.V.M., E.U., M.M.K, A.T., M.A. ; Writing – Original Draft, C.V.M.; Writing – Review & Editing, C.V.M., A.V.M., E.U., K.B.S., M.M.K, A.T., M.A., E.H.A, N.A., K.D.H., R.N., J.H.E., and C.H.C.; Funding Acquisition, C.H.C.; Resources, R.N. and C.H.C.; Supervision, K.D.H., R.N., J.H.E., and C.H.C.

## References

1 Bernardo, M. P. et al. PLA/Hydroxyapatite scaffolds exhibit in vitro immunological inertness and promote robust osteogenic differentiation of human mesenchymal stem cells without osteogenic stimuli. Scientific reports 12, 1–15 (2022).

2 Begum, S. A., Krishnan, P. S. G. & Kanny, K. Properties of poly (lactic Acid)/hydroxyapatite biocomposites for 3D printing feedstock material. Journal of Thermoplastic Composite Materials, 08927057231182165 (2023).

3 Huolman, R. & Ashammakhi, N. New multifunctional anti-osteolytic releasing bioabsorbable implant. Journal of Craniofacial Surgery 18, 295–301 (2007).

4 Länsman, S. et al. Poly-L/D-lactide (PLDLA) 96/4 fibrous implants: histological evaluation in the subcutis of experimental design. Journal of Craniofacial Surgery 17, 1121–1128 (2006).

5 Ashammakhi, N., Suuronen, R., Tiainen, J., Törmälä, P. & Waris, T. Spotlight on naturally absorbable osteofixation devices. Journal of Craniofacial Surgery 14, 247–259 (2003).

6 Zhang, Y. & Rehmmann, L. in Innovative and Emerging Technologies in the Bio-Marine Food Sector 417–439 (Elsevier, 2022).

7 Battafarano, G. et al. Strategies for bone regeneration: from graft to tissue engineering. International journal of molecular sciences 22, 1128 (2021).

8 Xu, Y. et al. Immunology and bioinformatics analysis of injectable organic/inorganic microfluidic microspheres for promoting bone repair. Biomaterials 288, 121685 (2022).

9 Xue, H. et al. Enhanced tissue regeneration through immunomodulation of angiogenesis and osteogenesis with a multifaceted nanohybrid modified bioactive scaffold. Bioactive Materials (2022).

10 Ma, B. et al. Hydroxyapatite nanobelt/polylactic acid Janus membrane with osteoinduction/barrier dual functions for precise bone defect repair. Acta Biomaterialia 71, 108–117 (2018).

11 Marycz, K. et al. Three dimensional (3D) printed polylactic acid with nano-hydroxyapatite doped with europium (III) ions (nHAp/PLLA@ Eu3+) composite for osteochondral defect regeneration and theranostics. Materials Science and Engineering: C 110, 110634 (2020).

12 Verardi, S., Lombardi, T. & Stacchi, C. Clinical and radiographic evaluation of nanohydroxyapatite powder in combination with polylactic acid/polyglycolic acid copolymer as bone replacement graft in the surgical treatment of intrabony periodontal defects: A retrospective case series study. Materials 13, 269 (2020).

13 Velard, F. et al. The effect of zinc on hydroxyapatite-mediated activation of human polymorphonuclear neutrophils and bone implant-associated acute inflammation. Biomaterials 31, 2001–2009 (2010).

14 Hunt, J. A. & Callaghan, J. T. Polymer-hydroxyapatite composite versus polymer interference screws in anterior cruciate ligament reconstruction in a large animal model. Knee Surgery, Sports Traumatology, Arthroscopy 16, 655–660 (2008).

15 Grandjean-Laquerriere, A. et al. Involvement of toll-like receptor 4 in the inflammatory reaction induced by hydroxyapatite particles. Biomaterials 28, 400–404 (2007).

16 Maduka, C. V. et al. Elevated oxidative phosphorylation is critical for immune cell activation by polyethylene wear particles. Journal of Immunology and Regenerative Medicine, 100069 (2023).

17 Maduka, C. V. et al. Glycolytic reprogramming underlies immune cell activation by polyethylene wear particles. Biomaterials Advances, 213495 (2023).

18 Li, X. et al. Glycolytic reprogramming in macrophages and MSCs during inflammation. Frontiers in Immunology 14 (2023).

19 Teissier, V. et al. Metabolic profile of mesenchymal stromal cells and macrophages in the presence of polyethylene particles in a 3D model. Stem Cell Research & Therapy 14, 1–16 (2023).

20 Agrawal, C. M. & Athanasiou, K. A. Technique to control pH in vicinity of biodegrading PLA-PGA implants. J Biomed Mater Res 38, 105–114, doi:10.1002/(sici)1097-4636(199722)38:2<105::aid-jbm4>3.0.co;2-u (1997).

21 Taylor, M. S., Daniels, A. U., Andriano, K. P. & Heller, J. Six bioabsorbable polymers: in vitro acute toxicity of accumulated degradation products. J Appl Biomater 5, 151–157, doi:10.1002/jab.770050208 (1994).

22 Maduka, C. V. et al. Polylactide degradation activates immune cells by metabolic reprogramming. Advanced Science, 2304632 (2023).

23 Maduka, C. V. et al. Stereochemistry Determines Immune Cellular Responses to Polylactide Implants. ACS Biomaterials Science & Engineering 9, 932–943 (2023).

24 Maduka, C. V. et al. Immunometabolic cues recompose and reprogram the microenvironment around biomaterials. bioRxiv, 2023.2007. 2030.551180 (2023).

25 Yang, L. et al. Macrophages at low-inflammatory status improved osteogenesis via autophagy regulation. Tissue Engineering Part A (2021).

26 Li, X. et al. Nanofiber-hydrogel composite–mediated angiogenesis for soft tissue reconstruction. Science translational medicine 11, eaau6210 (2019).

27 Edwards, F. C., Taheri, A., Dann, S. C. & Dye, J. F. Characterization of cytolytic neutrophil activation in vitro by amorphous hydrated calcium phosphate as a model of biomaterial inflammation. Journal of Biomedical Materials Research Part A 96, 552–565 (2011).

28 Velard, F. et al. Polymorphonuclear neutrophil response to hydroxyapatite particles, implication in acute inflammatory reaction. Acta Biomaterialia 5, 1708–1715 (2009).

29 Mahon, O. R. et al. Nano-particle mediated M2 macrophage polarization enhances bone formation and MSC osteogenesis in an IL-10 dependent manner. Biomaterials 239, 119833 (2020).

30 Boehler, R., Shin, S., Fast, A., Gower, R. M. & Shea, L. A PLG/HAp composite scaffold for lentivirus delivery. Biomaterials 34, 5431–5438 (2013).

31 Furukawa, T. et al. Bone bonding ability of a new biodegradable composite for internal fixation of bone fractures. Clinical Orthopaedics and Related Research® 379, 247–258 (2000).

32 Furukawa, T. et al. Histomorphometric study on high-strength hydroxyapatite/poly (Llactide) composite rods for internal fixation of bone fractures. Journal of Biomedical Materials Research: An Official Journal of The Society for Biomaterials, The Japanese Society for Biomaterials, and The Australian Society for Biomaterials and the Korean Society for Biomaterials 50, 410–419 (2000).

33 Ignatius, A. A., Betz, O., Augat, P. & Claes, L. E. In vivo investigations on composites made of resorbable ceramics and poly (lactide) used as bone graft substitutes. Journal of Biomedical Materials Research: An Official Journal of The Society for Biomaterials, The Japanese Society for Biomaterials, and The Australian Society for Biomaterials and the Korean Society for Biomaterials 58, 701–709 (2001).

34 Lebre, F. et al. The shape and size of hydroxyapatite particles dictate inflammatory responses following implantation. Scientific reports 7, 2922 (2017).

35 Shanley, L. C. et al. Macrophage metabolic profile is altered by hydroxyapatite particle size. Acta Biomaterialia 160, 311–321 (2023).

36 Pujari-Palmer, S. et al. In vivo and in vitro evaluation of hydroxyapatite nanoparticle morphology on the acute inflammatory response. Biomaterials 90, 1–11 (2016).

37 Ding, T. & Sun, J. Mechanistic understanding of cell recognition and immune reaction via CR1/CR3 by HAP-and SiO 2-NPs. BioMed Research International 2020 (2020).

38 Hua, Y. et al. Exposure to hydroxyapatite nanoparticles enhances Toll-like receptor 4 signal transduction and overcomes endotoxin tolerance in vitro and in vivo. Acta Biomaterialia 135, 650–662 (2021).

39 Graney, P. L., Roohani-Esfahani, S.-I., Zreiqat, H. & Spiller, K. L. In vitro response of macrophages to ceramic scaffolds used for bone regeneration. Journal of The Royal Society Interface 13, 20160346 (2016).

40 Mahon, O. R. et al. Orthopaedic implant materials drive M1 macrophage polarization in a spleen tyrosine kinase-and mitogen-activated protein kinase-dependent manner. Acta biomaterialia 65, 426–435 (2018).

41 Lange, T. et al. Proinflammatory and osteoclastogenic effects of betatricalciumphosphate and hydroxyapatite particles on human mononuclear cells in vitro. Biomaterials 30, 5312–5318 (2009).

42 Mahon, O. R., Kelly, D. J., McCarthy, G. & Dunne, A. Osteoarthritis-associated basic calcium phosphate crystals alter immune cell metabolism and promote M1 macrophage polarization. Osteoarthritis and Cartilage 28, 603–612 (2020).

43 Sadtler, K. et al. Developing a pro-regenerative biomaterial scaffold microenvironment requires T helper 2 cells. Science 352, 366–370 (2016).

44 Tannahill, G. et al. Succinate is a danger signal that induces IL-1β via HIF-1α. Nature 496, 238–242, doi:10.1038/nature11986 (2013).

45 Kauppinen, R. A., Sihra, T. S. & Nicholls, D. G. Aminooxyacetic acid inhibits the malate-aspartate shuttle in isolated nerve terminals and prevents the mitochondria from utilizing glycolytic substrates. Biochim Biophys Acta 930, 173–178, doi:10.1016/0167-4889(87)90029-2 (1987).

46 Jaynes, J. M. et al. Mannose receptor (CD206) activation in tumor-associated macrophages enhances adaptive and innate antitumor immune responses. Science translational medicine 12, eaax6337 (2020).

47 Niedermeier, M. et al. CD4+ T cells control the differentiation of Gr1+ monocytes into fibrocytes. Proceedings of the National Academy of Sciences 106, 17892–17897 (2009).

48 Werner, Y. et al. Cxcr4 distinguishes HSC-derived monocytes from microglia and reveals monocyte immune responses to experimental stroke. Nature neuroscience 23, 351–362 (2020).

49 Huang, L.-R. et al. Intrahepatic myeloid-cell aggregates enable local proliferation of CD8+ T cells and successful immunotherapy against chronic viral liver infection. Nature immunology 14, 574–583 (2013).

50 Sadtler, K. et al. Divergent immune responses to synthetic and biological scaffolds. Biomaterials 192, 405–415 (2019).

51 Guo, M. et al. In vivo immuno-reactivity analysis of the porous three-dimensional chitosan/SiO2 and chitosan/SiO2/hydroxyapatite hybrids. Journal of Biomedical Materials Research Part A 106, 1223–1235 (2018).

52 Jha, A. K. et al. Network integration of parallel metabolic and transcriptional data reveals metabolic modules that regulate macrophage polarization. Immunity 42, 419–430, doi:10.1016/j.immuni.2015.02.005 (2015).

53 Zhao, Q. et al. Dual-wavelength photosensitive nano-in-micro scaffold regulates innate and adaptive immune responses for osteogenesis. Nano-Micro Letters 13, 1–20 (2021).

54 Korangath, P. et al. Targeting Glutamine Metabolism in Breast Cancer with AminooxyacetateTargeting Glutamine Metabolism in Breast Cancer. Clinical cancer research 21, 3263–3273 (2015).

